# pGGTOX: a versatile plasmid platform for recombination-based genome editing across *Enterobacteriaceae*

**DOI:** 10.1101/2025.11.11.687829

**Authors:** Min Liu, Enyuan Tian, Xiaodi Cui, Ke Liu, Liya Feng, Yuheng Liu, Yujiao Wang, Xiaohong Shi, Liang Chen, Mingju Hao

## Abstract

Precise genome editing in *Enterobacteriaceae* is essential for studying gene function, pathogenesis, and antimicrobial resistance, yet many current systems face host-specific and efficiency limitations. We developed pGGTOX, a modular plasmid platform that enables efficient homologous recombination–mediated genome editing across diverse *Enterobacteriaceae*, including *Escherichia coli*, *Klebsiella pneumoniae*, *Salmonella enterica*, and *Enterobacter intestinihominis*. The system integrates a rhamnose-inducible toxin (MqsR) for stringent counterselection, a sfGFP reporter for visual tracking of recombination events, Golden Gate cloning for rapid assembly of homologous arms, an FRT-flanked resistance cassette for marker removal, and an *oriT* sequence for conjugative transfer. Together with the companion plasmid pCP20-oriT, pGGTOX supports precise, marker-free genomic modification. Using pGGTOX, we achieved targeted deletions of *dapA* in *E. coli* and *mrkCD* in carbapenem-resistant *K. pneumoniae*, both with 100% efficiency. The *dapA* mutant exhibited diaminopimelate auxotrophy, while *mrkCD* deletion markedly reduced biofilm formation, consistent with the loss of function associated with these genes. pGGTOX also enabled deletion of a 43.1-kb type IV secretion gene cluster (*tra*) from an IncN/FII plasmid in *E. intestinihominis* and insertion of a 10-kb CRISPR–Cas9 plasmid-curing module (pCasCure) into an *S. enterica* IncX1 plasmid. Deletion of the *tra* gene cluster resulted in a substantial reduction in plasmid conjugation efficiency. Conjugative transfer of the engineered IncX1-pCasCure plasmid into *K. pneumoniae* facilitated CRISPR-mediated curing of *bla*_KPC_, sensitizing carbapenem resistance to susceptibility. In summary, pGGTOX provides a versatile, efficient, and broadly applicable platform for genome engineering and CRISPR delivery in *Enterobacteriaceae*, expanding the toolkit for bacterial genetics and translational antimicrobial research.

**Importance:** Precise genetic manipulation in *Enterobacteriaceae* remains a major technical challenge, particularly for non-model or multidrug-resistant strains. We developed pGGTOX, a versatile and broadly applicable plasmid platform that enables efficient, marker-free genome editing through homologous recombination. By integrating stringent counterselection, visual screening, modular cloning, and conjugative transfer, pGGTOX simplifies construction and streamlines editing across multiple clinically relevant species. We demonstrate its utility in deleting chromosomal and plasmid-borne loci, inserting large genetic modules, and delivering CRISPR–Cas9 systems for targeted elimination of antibiotic resistance genes. This platform expands the molecular toolkit for functional genomics and provides a powerful new strategy for dissecting bacterial virulence, resistance, and plasmid biology.

## Introduction

Bacterial genome editing is a powerful approach for dissecting gene function, exploring pathogenic mechanisms, and enabling metabolic engineering in both laboratory and applied settings(1). Within the *Enterobacteriaceae*—including *Escherichia coli*, *Klebsiella pneumoniae*, and *Salmonella enterica*—precise genetic manipulation has broad implications for basic microbiology, biotechnology, and public health. Over the years, editing methods have advanced from allelic exchange using suicide plasmids to phage-derived recombination systems such as the λ-Red recombinase(2–4), which allows efficient integration of linear DNA fragments with short homology arms. This system, commonly referred to as “recombineering,” has become a cornerstone of bacterial genetics due to its versatility and compatibility across multiple species.

More recent innovations have combined homologous recombination with CRISPR– Cas counterselection to improve editing efficiency and stringency(5, 6). By introducing targeted double-stranded breaks, CRISPR–Cas strongly enriches for recombination events, and in *E. coli* and *K. pneumoniae*, this strategy has markedly accelerated genome editing(5, 7). However, CRISPR-based systems face notable limitations: delivery of large editing complexes can be inefficient, double-strand breaks can be cytotoxic, and species-specific barriers often reduce efficacy in non-model or clinically relevant strains(8, 9). These challenges restrict the routine use of CRISPR in many bacteria, underscoring the continued importance of recombination-based tools.

In this study, we describe the development of pGGTOX, a plasmid-based platform for homologous recombination–mediated genome editing in *Enterobacteriaceae*. pGGTOX integrates key design features to improve precision, flexibility, and ease of use, including a rhamnose-inducible toxin for stringent negative selection(10–12), Golden Gate cloning for seamless insertion of homologous arms(13), an FRT-flanked resistance marker for removable selection(14), a GFP-based counterselection system for visual screening, and an *oriT* sequence for conjugative transfer. By combining stringent selection, modular cloning, and broad host applicability, pGGTOX provides a practical and efficient solution for complex genetic manipulations and expands the toolkit available for genome editing in diverse *Enterobacteriaceae*.

## Results

### Construction of pGGTOX and pCP20-oriT

The plasmids pGGTOX and pCP20-oriT were constructed to facilitate gene knockout and the subsequent removal of antibiotic resistance marker. pGGTOX was designed by integrating essential genetic elements from the plasmids pTOX, pGGA-select, and pSZU941-GFP. The *mqsR* toxin from pTOX ensures effective counterselection, minimizing the likelihood of escape events (10). Additionally, the sfGFP fluorescence module enables visualization of single and double crossover events, aiding in the monitoring of recombination success. The *oriT* element enables the conjugative transfer of pGGTOX, while the R6K origin of replication restricts its replication to host strains expressing the *pir* gene. The apramycin resistance gene, *aac3(IV),* was incorporated to enable its use in clinically drug resistant hosts, as most MDR enterobacterial strains remain susceptible to apramycin (15) (Fig. 1A). Furthermore, alternative antimicrobial selection markers can be readily incorporated during pGGTOX assembly (see Methods for details). To facilitate insertion of the homologous fragments, pGGTOX incorporates two Type IIS restriction sites (BsaI/BsmB1). Upon enzymatic digestion (BsaI or BsmB1), the plasmid is cleaved into two equimolar fragments, each carrying pre-designed 4-base overhangs that guide precise homologous assembly.

**Fig. 1.**
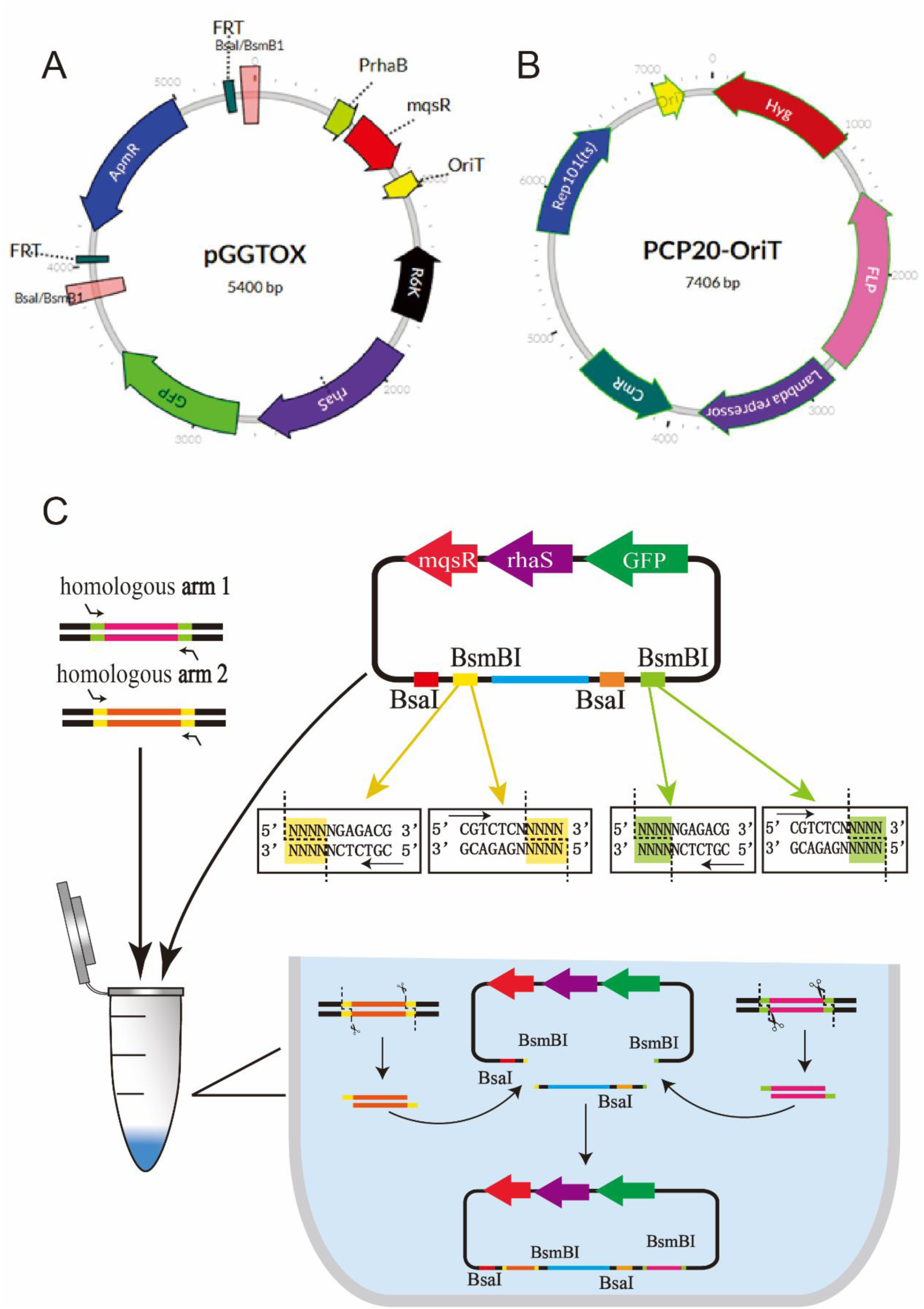
Plasmid map of pGGTOX, pCP20-oriT and one-step Golden Gate assembly of homologous fragments into pGGTOX. (A) pGGTOX: R6K ori, R6K origin of replication; mobRP4, mobilization region from the RP4 conjugative plasmid; rhaS, rhamnose transcriptional activator gene; ApmR, apramycin resistance cassette; pRha, rhamnose promoter. GFP, green fluorescent protein encoding gene; mqsR, mqsR toxin encoding gene; The vertical bars of various widths represent type II restriction recognition sites BsaI and BsmBI. (B) PCP20-oriT: HygR, hygromycin resistance cassette; FLP, Flp recombinase; CmR, Chloramphenicol resistance. (C) Homologous fragments were amplified by PCR with flanking Type IIS restriction sites to enable directional cloning. The purified PCR products were combined with the pGGTOX vector, a type IIS restriction enzyme (BsmB1 or BsaI), T4 DNA ligase, and appropriate buffer in a single-tube Golden Gate reaction. Each fragment contained designed 4-base overhangs to guide precise ligation. Following assembly, the reaction mixture was used to transform chemically competent *E. coli* WM3064 cells for plasmid recovery. The plasmid maps were generated with Angular Plasmid.

Similarly, the pCP20-oriT plasmid was constructed by incorporating essential fragments from pCP20, pSGKp-hygromycin and pdCas-J23119-RFP-oriT plasmids, ensuring the presence of sfGFP, *oriT,* FLP recombinase and resistance markers (Fig. 1B). These modifications facilitate transference of pCP20-oriT into recipient cells via conjugation, monitoring conjugates and expand the resistant makers of the initial pCP20 plasmid.

### Genomic editing workflow using pGGTOX

#### One-Step Golden Gate assembly of homologous fragments into pGGTOX

Homologous DNA inserts were generated via PCR using primers designed to incorporate Type IIS restriction sites and 4-base pair overhangs, enabling directional and sequential assembly. The purified PCR products were combined with the pGGTOX vector, a Type IIS restriction enzyme (BsmBI or BsaI), T4 DNA ligase, and appropriate reaction buffer in a single-tube Golden Gate assembly. Thermocycling between digestion and ligation phases facilitated seamless junction formation, guided by the engineered overhangs. An apramycin resistance cassette was strategically placed between the homologous arms to allow for selection of recombinant clones (Fig. 1C).

#### Induction of homologous recombination for targeted gene editing

To initiate genome editing, assembled pGGTOX constructs were first introduced into chemically competent *E. coli* WM3064 cells. This strain is auxotrophic for diaminopimelate (DAPA), allowing for selective growth and facilitating identification of successful transconjugants. Colony PCR targeting the junction regions was performed to confirm correct integration of all fragments into the pGGTOX backbone (Fig. S1).

Verified recombinant plasmids were subsequently transferred into target bacterial strains via conjugation. Following conjugation, homologous recombination could proceed either through a single crossover at one homologous arm or via a double crossover involving both arms (Fig. 2A).

**Fig. 2.**
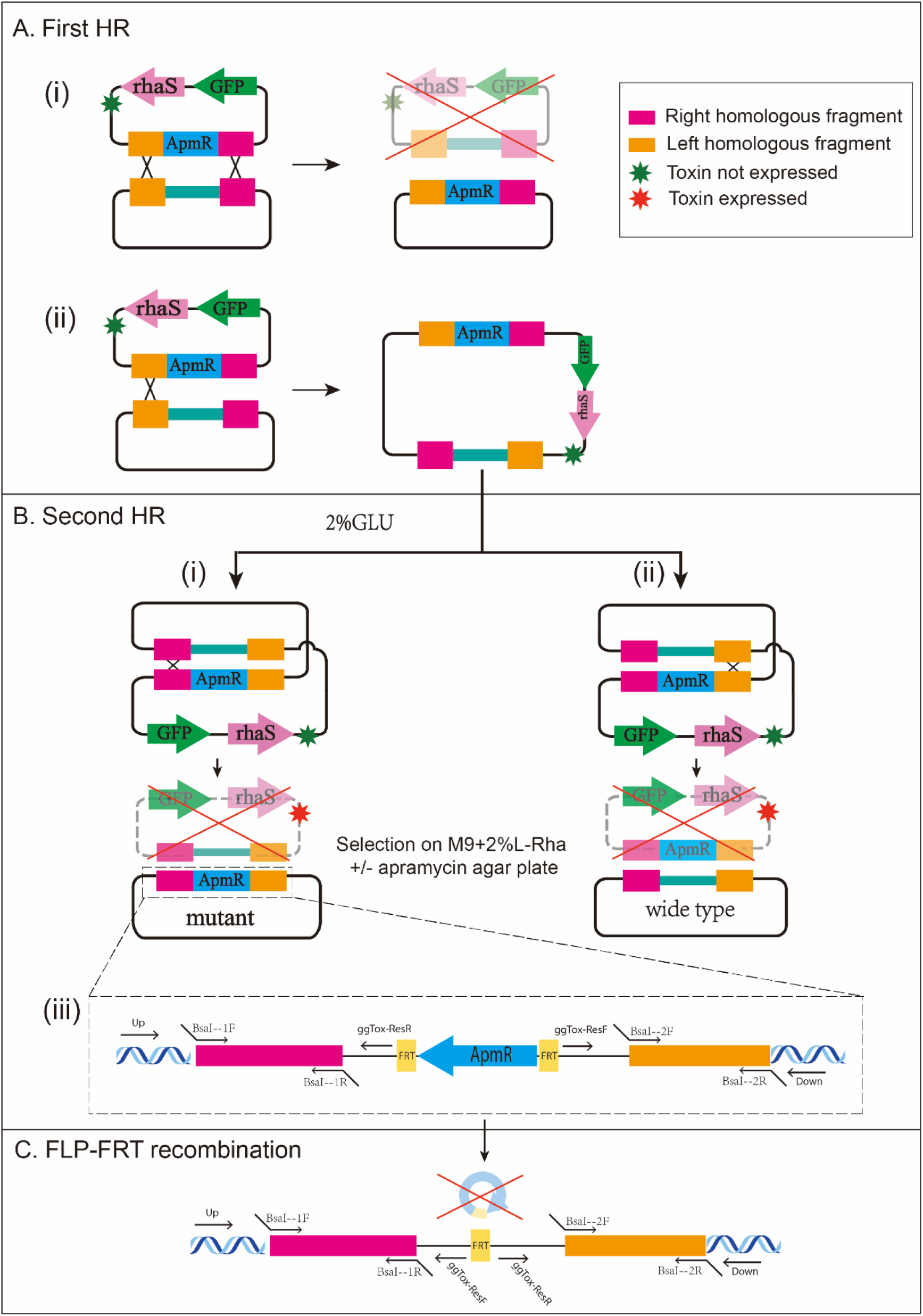
Genomic editing workflow using pGGTOX. (A) Homologous fragments flanking the target gene were inserted into pGGTOX and introduced into the host strain to initiate the first homologous recombination event. (i) Successful recombination at both flanking regions results in precise genome editing. (ii) Merodiploid colonies containing the integrated plasmid can be identified by GFP fluorescence. (B) The second recombination event was induced by culturing merodiploid strains in medium supplemented with 2% glucose, followed by counter-selection using rhamnose-induced expression of the mqsR toxin. (i) Strains that undergo recombination at the second homologous site retain the desired genomic edit and survive on apramycin-supplemented plates. (ii) Strains that revert to the wild-type sequence are eliminated due to apramycin sensitivity. (iii) Primer pairs (Up/ggTox-ResR and Down/ggTox-ResF) used for PCR confirmation of editing events and their binding sites are illustrated at the bottom of the panel. (C) Following confirmation of successful genome editing, the antibiotic resistance marker can be excised via FLP recombinase-mediated recombination at flanking FRT sites.

Initial single-arm recombination was monitored using GFP fluorescence, which served as a visual marker for integration. Colonies exhibiting GFP signal were selected for a second recombination event at the opposite homologous arm (Fig. 2B). Completion of both recombination steps resulted in precise genomic modification, confirming the effectiveness of the pGGTOX system for targeted gene editing (Fig. 2B).

#### Removal of antibiotic resistance genes via pCP20-oriT

The pCP20-oriT plasmid was used to facilitate the excision of apramycin resistance cassettes via FLP-mediated recombination (Fig. 2C). Conjugation into recombined bacterial strains resulted in the excision of the FRT-flanked resistance gene and the pCP20-oriT helper plasmid simultaneously. The target strains restored susceptibility to apramycin. The effectiveness of this approach demonstrates the utility of the pCP20-oriT system for marker excision in genome editing applications.

### Targeted knockout of *dapA* and *mrkCD* in *Enterobacteriaceae* using pGGTOX

To evaluate the utility of the pGGTOX vector system for gene deletion in *Enterobacteriaceae*, we selected *dapA* in *E. coli* Nissle 1917 and *mrkCD* in carbapenem-resistant *K. pneumoniae* JNQH373 for gene knockout. The *dapA* gene encodes dihydrodipicolinate synthase, a key enzyme in the diaminopimelate (DAP) pathway essential for peptidoglycan synthesis and lysine biosynthesis(16). Elimination of *dapA* can lead to auxotrophic phenotype. The *mrkABCDF* gene cluster encodes type 3 fimbriae, which mediate biofilm formation and surface adhesion in *K. pneumoniae* (*17*). Elimination of *mrkABCDF* can lead to decreased biofilm formation. Homology arms flanking each target locus were PCR-amplified and directionally assembled into the pGGTOX backbone using one-step Golden Gate cloning. The resulting constructs, pGGTOX-*dapA* and pGGTOX-*mrkCD*, were verified by colony PCR and introduced into recipient strains via conjugation. Green fluorescent colonies were observed post-conjugation, indicating successful single-crossover integration and formation of merodiploid intermediates. To resolve the second crossover, merodiploids were plated on M9 minimal agar supplemented with rhamnose to induce toxin expression and counter-select against cells retaining the integrated plasmid. Only cells that had undergone a successful double-crossover event survived, forming non-fluorescent (white) colonies. PCR screening of seven randomly selected *dapA* knockout candidates and four *mrkCD* knockout candidates confirmed precise deletions in all cases (Fig. 3A-B), demonstrating the efficiency and reliability of the pGGTOX system for genome editing in both *E. coli* and *K. pneumoniae*. To finalize the editing process, the apramycin resistance marker was excised using FLP recombinase-mediated recombination at flanking FRT sites, thereby restoring a marker-free genomic locus (Fig. 3C).

**Fig. 3.**
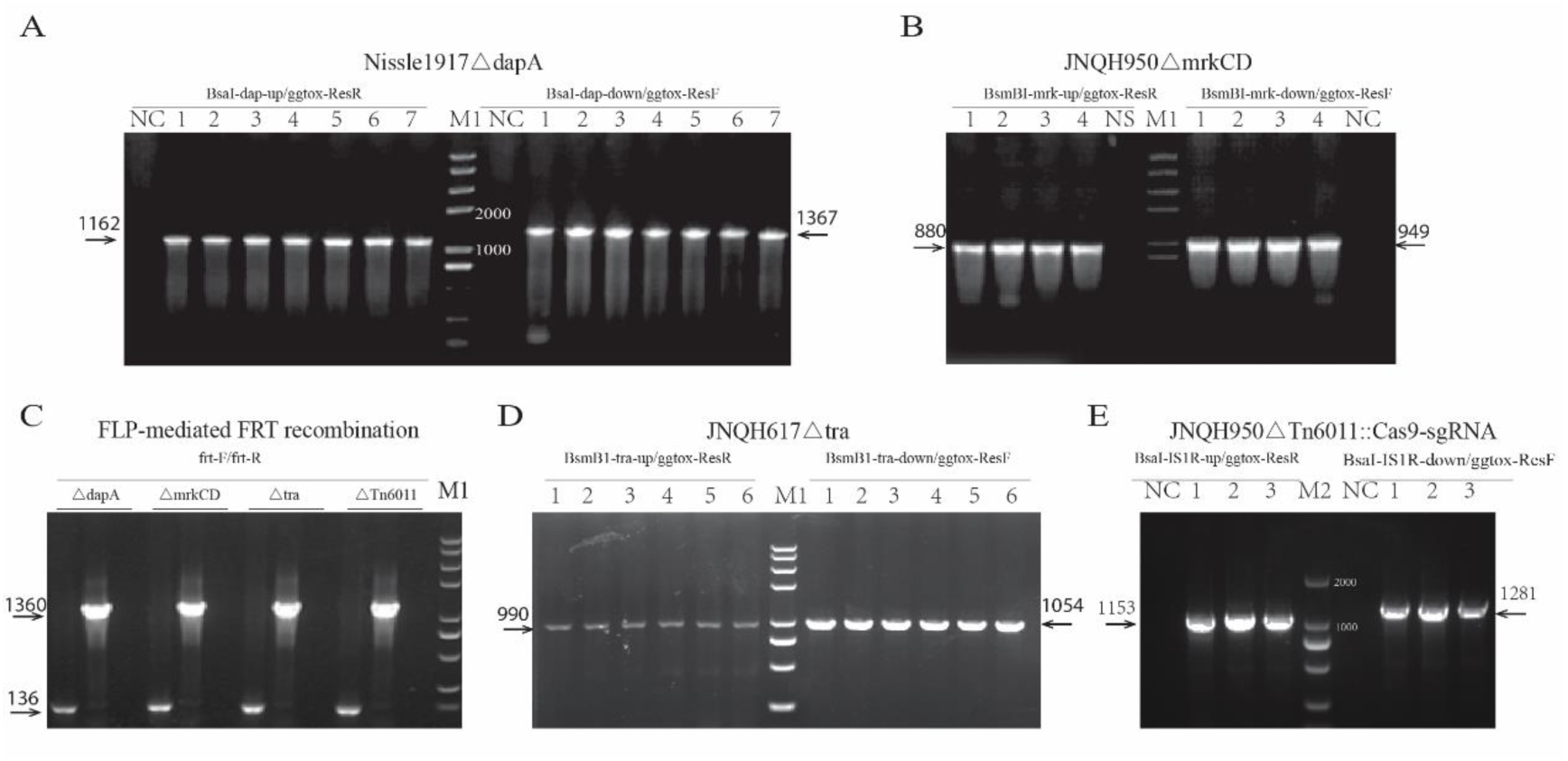
Validation of targeted gene deletions by PCR. (A) Deletion of the *dapA* gene in *E. coli s*train Nissle 1917. (B) Deletion of *mrkCD* in *S.enterica* strain JNQH950. (C) FLP-mediated FRT recombination to excise apramycin resistance genes. For each mutant strain, gel lanes (left to right) represent colonies that underwent FRT recombination, colonies that did not undergo recombination, negative control. (D) Deletion of the *tra* cassette in *E. intestinihominis* JNQH617. (E) Replacement of *Tn*6011 with Cas9/sgRNA in *S. enterica* JNQH950. Primer pairs used for amplification are indicated above each gel panel. Expected sizes of PCR products are shown on either the left or right side of each gel image. M1: DNA ladder (from top to bottom, the sizes of the bands are 8000, 5000, 3000, 2000, 1000, 750, 500, 250 and 100 bp); M2: DNA ladder (from top to bottom, the sizes of the bands are 2000, 1000, 750, 500, 250, 100 bp).

### Targeted knockout of a 43.1 kb *tra* gene cluster using pGGTOX

The *tra* gene cluster is a key component of conjugative plasmids, mediating the horizontal transfer of genetic material between bacterial cells. It typically encodes the Type IV coupling protein (T4CP), the Type IV secretion system (T4SS), and a relaxase enzyme—each playing a coordinated role in the conjugation process (18–20).

To investigate the role of the *tra* cluster in conjugation efficiency, we targeted a 43.1 kb *tra* locus located on a hybrid IncN/FII plasmid in *E. intestinihominis* strain JNQH617. Homology arms flanking the *tra* region were PCR-amplified and assembled into the pGGTOX backbone using Golden Gate cloning. The resulting construct, pGGTOX-*tra*, was introduced into JNQH617 via conjugation, with *E. coli* WM3064 serving as the donor strain.

Following conjugation, only non-fluorescent (white) colonies were recovered on apramycin-selective agar, indicating potential recombination events. Six colonies were randomly selected for PCR validation, all of which confirmed successful deletion of the entire *tra* cluster. These results suggest that homologous recombination occurred efficiently in a single step, without the need for a secondary crossover event (Fig. 3D).

### Insertion of pCasCure *Cas9-sgRNA*_KPC_ module into the IncX1 plasmid using pGGTOX-Cas9-sgRNA_KPC_ plasmid

We previously developed a CRISPR-Cas9-based system, pCasCure, which demonstrated effective curing of carbapenemase genes and resistance plasmids, thereby resensitizing carbapenem-resistant *Enterobacteriaceae* (CRE) to carbapenem antibiotics(21). However, a key limitation of pCasCure is its reliance on electroporation for delivery, which restricts its practical application, particularly in clinical or environmental settings.

To overcome this barrier, we explored the use of the IncX1 plasmid as a natural delivery vehicle. IncX1 plasmids are widely distributed among *Enterobacteriaceae* and exhibit high conjugative transfer efficiency(22, 23), making them ideal candidates for disseminating genome editing tools. Although IncX1 plasmids typically lack antibiotic resistance genes, they often carry the Tn*6011* transposon, which encodes the *mrkABCDF* virulence cluster associated with biofilm formation(22, 23).To construct a mobile version of the pCasCure system, we used pGGTOX to insert the Cas9-sgRNA_KPC_ module into the IncX1 plasmid of *S. enterica* strain JNQH950, replacing the Tn*6011* transposon. This strategy enabled the development of a conjugation-capable, self-propagating CRISPR/Cas9 platform for targeted elimination of resistance genes, offering a promising alternative to electroporation-based delivery methods.

To enable CRISPR-mediated targeting of *bla*_KPC_, we successfully constructed the recombinant plasmid pGGTOX-Cas9-sgRNA_KPC_. The Cas9 coding sequence and sgRNA-N20_KPC_ element were amplified from the pCasCure-N20_KPC_(21) template and assembled into a linearized pGGTOX backbone using In-Fusion cloning. The resulting plasmid supports targeted delivery of *Cas9* and sgRNA into recipient strains, providing a platform for CRISPR-based interference against carbapenem resistance determinants (Fig. 4A).

**Fig. 4.**
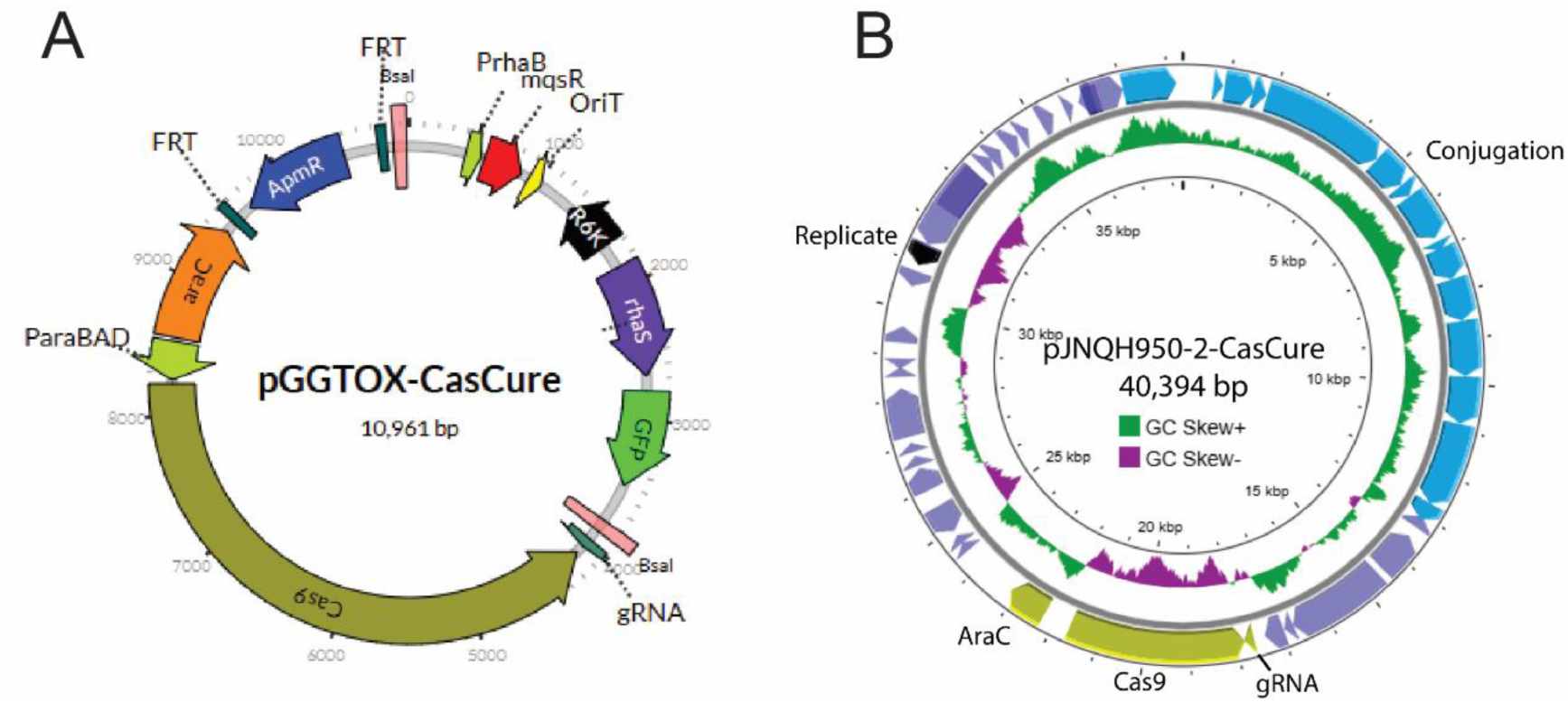
Plasmid map of pGGTOX-CasCure and pJNQH950-2-CasCure. (A) pGGTOX-CasCure: rhaS, Rhamnose system transcriptional activator; Apmr, apramycin resistance gene *aac(3’)-IV*; araC, arabinose operon regulatory protein. ParaBAD, promoter of the arabinose operon. (B) pJNQH950-2-CasCure: The self-conjugative pJNQH950-2-CasCure plasmid was constructed by insertion of CRISPR/Cas9 system into a highly conjugative IncX1 plasmid substituting the *Tn6011* transposon. The conjugation elements, CRISPR/Cas9, replicon and other coding sequences are denoted in different colors. The plasmid maps were generated with Angular Plasmid (A) and Proksee (B).

Homology arms flanking the native Tn*6011* transposon were PCR-amplified and assembled into the pGGTOX-Cas9-sgRNA_KPC_ backbone. The resulting construct was mobilized into JNQH950 via conjugation. As observed in previous large-locus knockouts, recombination resolved directly at both ends of the homologous region without requiring a secondary crossover event (Fig. 2Ai). Colony PCR confirmed successful replacement of Tn*6011* with the Cas9-sgRNA_KPC_ module in all three tested colonies (Fig. 3E), generating the engineered plasmid pJNQH950-2-CasCure_KPC_ (Fig. 4B). This modified IncX1 plasmid was subsequently transferred into strain JNQH373 via conjugation to enable CRISPR-mediated curing of the *bla*_KPC_ resistance gene.

### Phenotypic consequences of targeted gene disruption and CRISPR-Cas9 delivery

Deletion of the *dapA* gene in *E. coli* Nissle 1917 resulted in a clear auxotrophic phenotype. The mutant strain failed to grow in diaminopimelate (DAP)-deficient media but regained normal growth upon DAP supplementation (Fig. 5A), confirming both the successful knockout and its functional impact on cell wall biosynthesis.

**Fig. 5.**
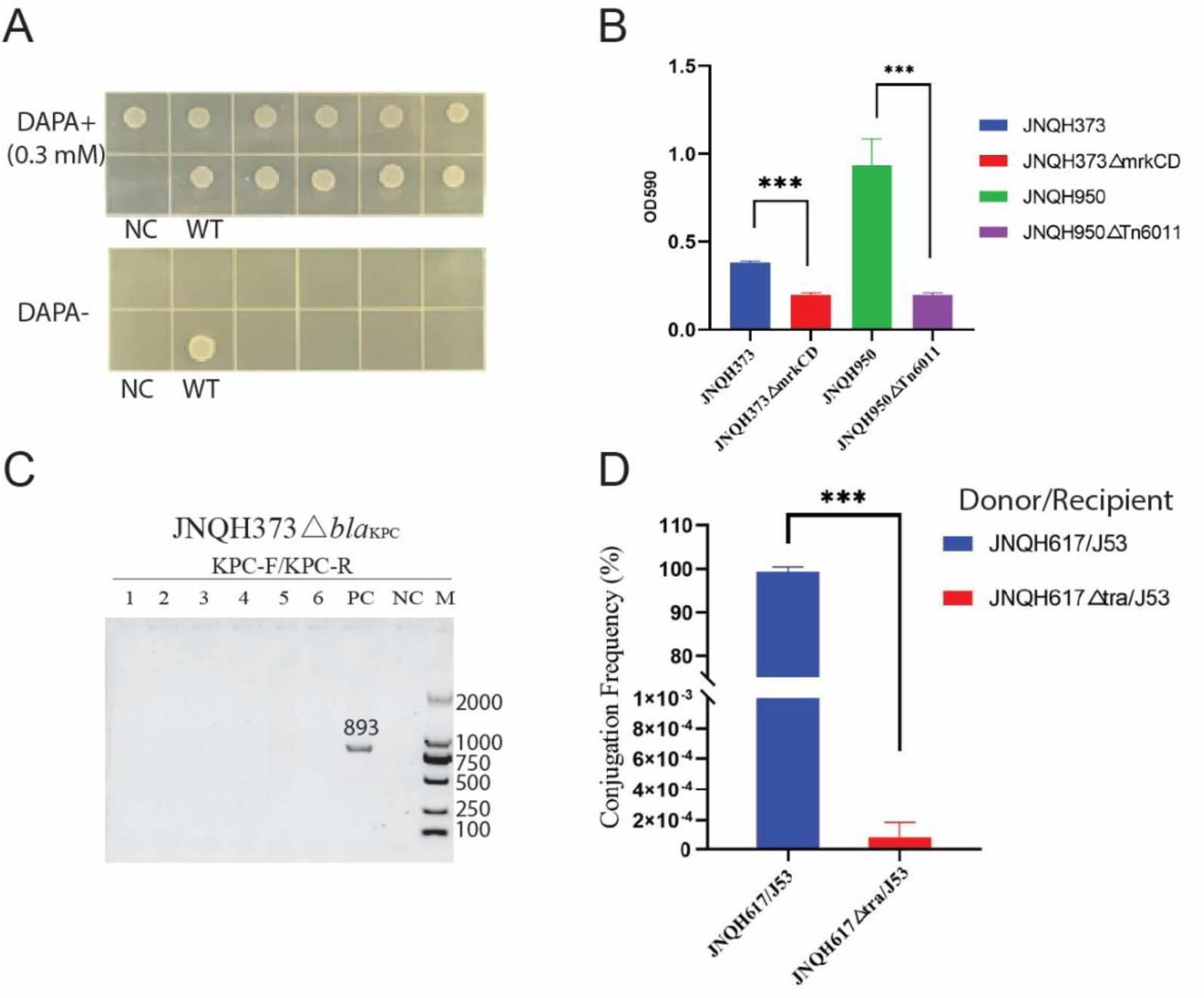
Phenotypic consequences of targeted gene disruption and CRISPR-Cas9 mediated *bla*_KPC_ curing. (A) The *dapA* knockout resulted in DAPA auxotrophy, with growth restored upon DAPA supplementation. (B) Deletion of *mrkCD* in *K. pneumoniae* and Tn*6011* harboring *mrkABCDF* gene loci significantly decreased biofilm formation. (C) The engineered IncX1 plasmid carrying Cas9-sgRNA_KPC_ resulted in efficient *bla*_KPC_ curing in carbapenem resistant *K. pneumoniae* via conjugation. (D) Deletion of the *tra* gene cluster in *E. intestinihominis* JNQH617 reduce the conjugation efficiency of the IncN/FII plasmid significantly. NC, negative control; WT, wild type *E.coli* Nissil 1917; \*\*\**P* < 0.001

In *K. pneumoniae* strain JNQH373, targeted deletion of *mrkCD* led to a substantial reduction in biofilm formation (Fig. 5B), underscoring the essential role of the *mrkABCDF* gene cluster in surface adhesion and biofilm development. Similarly, replacement of the Tn*6011* transposon—harboring *mrkABCDF*—with the Cas9-sgRNA_KPC_ module in *S. enterica* strain JNQH950 significantly impaired biofilm formation (Fig. 5B).

The engineered IncX1 plasmid carrying Cas9-sgRNA_KPC_ was successfully conjugated from JNQH950 into carbapenem-resistant strain JNQH373. Upon arabinose induction, CRISPR-mediated targeting of *bla*_KPC_ resulted in efficient gene elimination (Fig. 5C), as evidenced by a marked reduction in minimum inhibitory concentrations (MICs) of imipenem and meropenem from >8 μg/mL to <0.25 μg/mL. These results demonstrate the potential of conjugation-based delivery of CRISPR-Cas9 systems for precise elimination of antibiotic resistance determinants.

In addition, deletion of the *tra* gene cluster in *E. intestinihominis* strain JNQH617 resulted in a marked reduction in the conjugation efficiency of the IncN/FII plasmid (Fig. 5D). This finding underscores the critical role of the *tra* locus in facilitating plasmid-mediated gene transfer and highlights its contribution to the dissemination of antibiotic resistance within *Enterobacteriaceae*.

## Materials and Methods

### Bacterial strains and plasmids

Strains and plasmids are detailed in Table 1. The bacterial strains used in this study included *E. coli* WM3064, *E. coli* Nissle 1917, *K. pneumoniae* JNQH373, *E. intestinihominis* JNQH617 and Nontyphoid *Salmonella* serotype Thompson JNQH950. WM3064 was used as a donor strain for conjugation experiments. *K. pneumoniae* JNQH373 carried the *mrkABCDF* type 3 fimbriae in the chromsome, while *Salmonella* JNQH950 harbored the *mrkABCDF* gene cluster carried by *Tn*6011 on the IncX1 plasmid(**24**). Strains were routinely grown at 37℃ for up to 18 h in Luria broth Miller (LB) or on LB agar medium supplemented with antibiotics when needed. Antibiotics were used at the following working concentrations: apramycin 30 µg/ml, chloramphenicol 25 µg/ml, ampicillin 100 µg/ml, kanamycin 50 µg/ml, streptomycin 50 µg/ml, DAP auxotrophy was complemented by adding DAP at a final concentration of 0.3 mM.

**Table 1.**
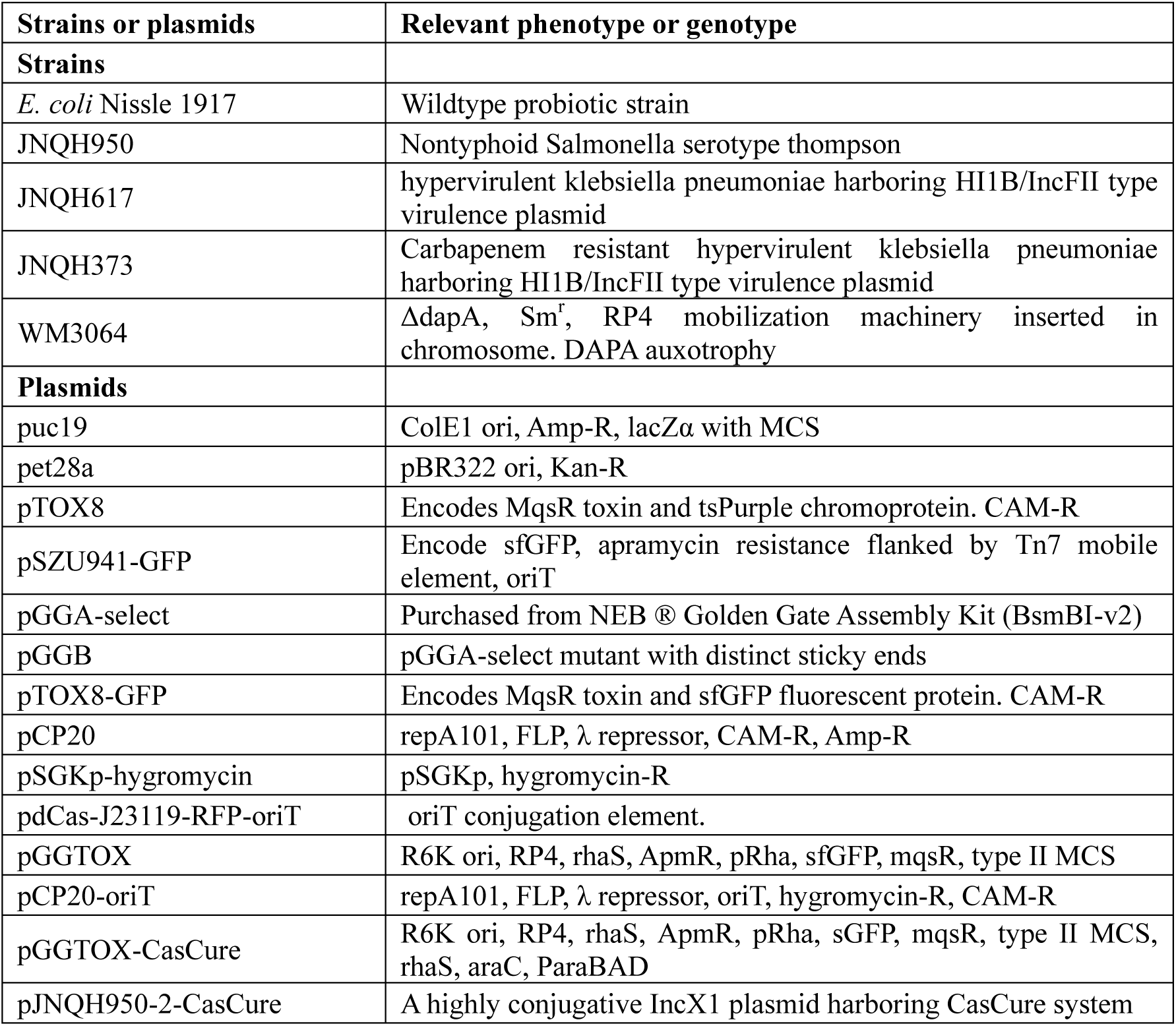
Strains or plasmids used in this study.

pSZU941-GFP was a gift from Yuzhang He. pSGKp-hygromycin was a gift from Yingying Hao. pGGAselect was purchased from New England Biolabs as a component of NEB® Golden Gate Assembly Kit (NEB #E1602S/L); pTOX8 was a gift from Matthew Waldor (Addgene plasmid #127453; http://n2t.net/addgene:127453; RRID:Addgene_127453); pdCas-J23119-RFP-oriT was constructed in our lab previously.

### Construction of pGGTOX plasmid

The primers used for the construction of mutant strains are listed in Table 2. We first modified the original plasmid pGGA-select. The purpose of this modification was to introduce mutants to produce different sticky ends after *BsaI* and *BsmBI* digestion, ensuring the correct orientation of homologous arm insertion. Specifically, using pGGA-select as a template, we amplified the DNA fragment using the pGGE1/pGGE2 primer pair. The amplified fragment was phosphorylated and ligated to generate the new plasmid, which was then linearized using *NotI-HF* single digestion to serve as the backbone for pGGB plasmid. Next, we amplified a DNA fragment containing the original *BsaI* and *BsmBI* restriction sites from pGGA-select using the Bsa-BsmB-F/Bsa-BsmB-R primer pair. The apramycin resistance gene *aac(3’)-IV* was amplified from pSZU941-GFP using the pGG-Apr-F/pGG-Apr-R primer pair. These three DNA fragments were assembled using seamless cloning, forming a circularized plasmid of pGGB. Next, using the pGGB as a template, we amplified a DNA fragment containing *aac(3’)-IV* and multiple cloning sites using primers gg-Apr-ggF and gg-Apr-ggR to serve as the backbone for pGGTOX.

**Table 2.**
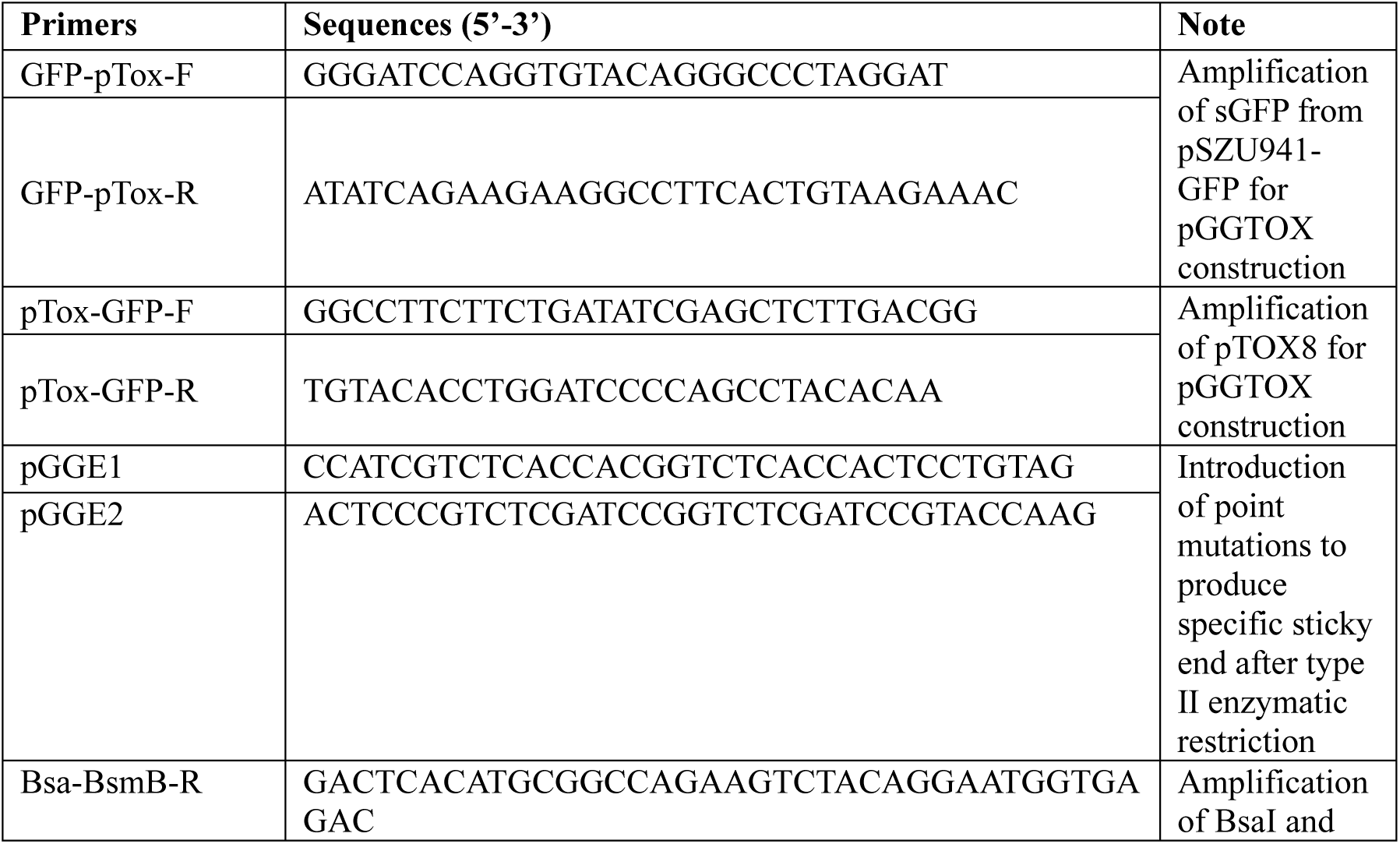
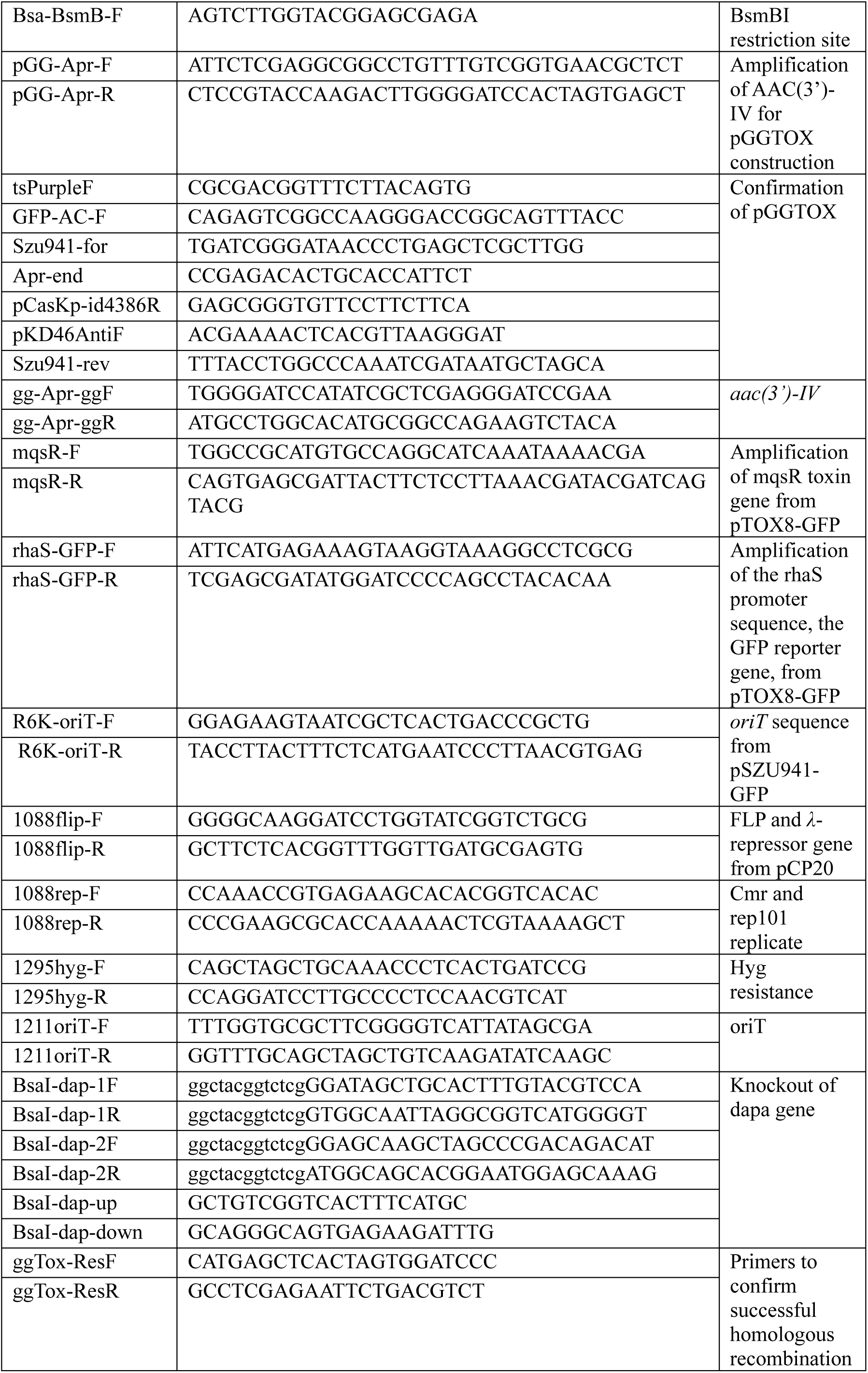
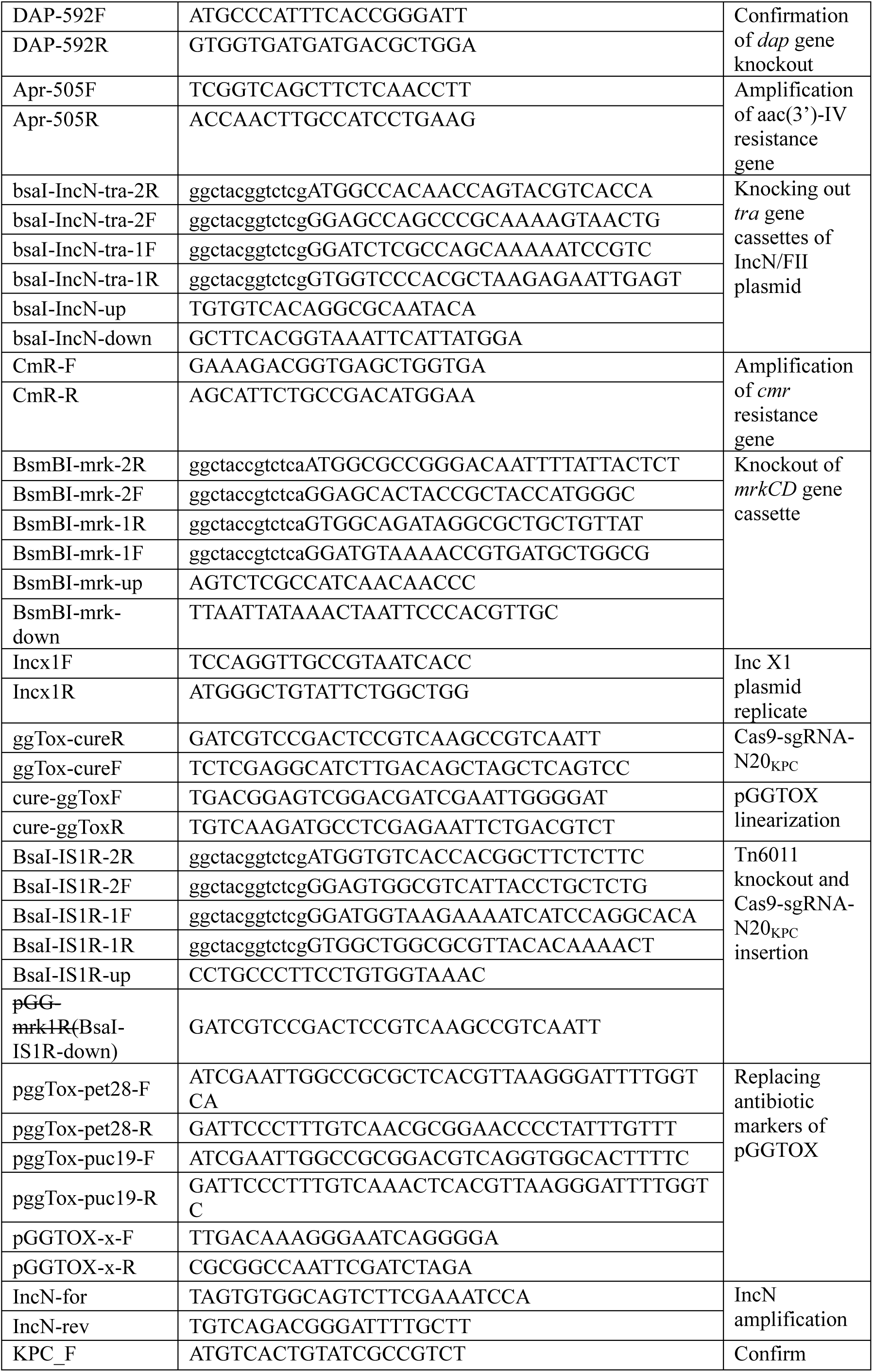
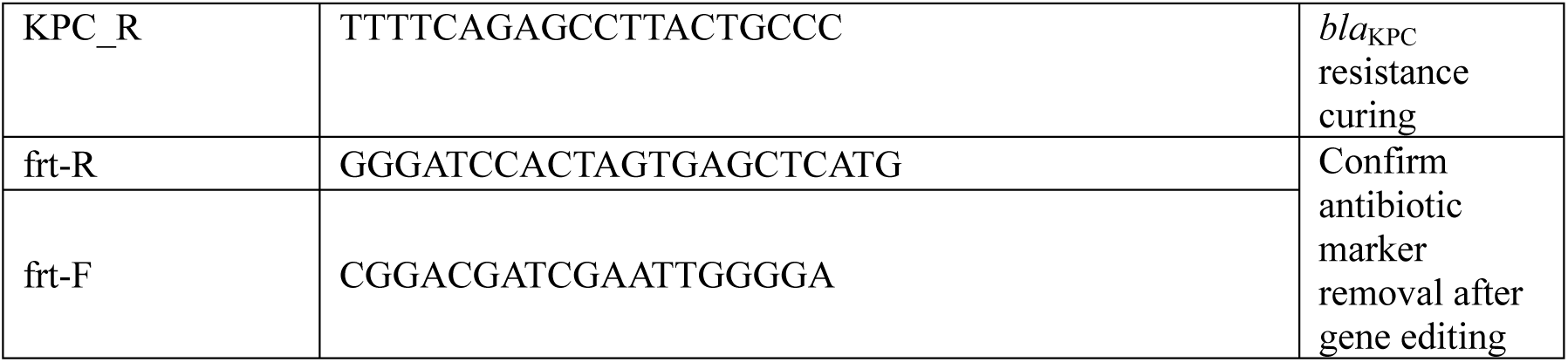
Primers used in this study.

Next, we designed and constructed pTOX8-GFP by replacing the tsPurple gene with the sGFP gene. First, we used plasmid pTOX8 as a template and designed primers pTox-GFP-F and pTox-GFP-R to amplify the plasmid backbone. Next, the sfGFP gene was amplified from plasmid pSZU941-GFP with primers GFP-pTox-F and GFP-pTox-R, serving as the insert fragment. The purified backbone and insert were then subjected to In-Fusion cloning, resulting in the pTOX8-GFP. Subsequently, we amplified the *rhaS* promoter sequence, the sfGFP reporter gene, and the *mqsR* toxin gene from pTOX8-GFP using primer pairs rhaS-GFP-F/rhaS-GFP-R and mqsR-F/mqsR-R. Additionally, the *oriT* sequence, essential for conjugative transfer, was amplified from pSZU941-GFP using primers R6K-oriT-F and R6K-oriT-R. These four DNA fragments were assembled into a single construct via seamless cloning. To confirm the accuracy of the assembled plasmid, we designed specific validation primers (tsPurpleF/GFP-AC-F, Szu941-for/Apr-end, Bsa-BsmB-F/pCasKp-d4386R, and pKD46AntiF/Szu941-rev). Further, the pGGTOX variants carrying ampicillin (pGGTOX-amp) or kanamycin (pGGTOX-kana) resistance markers were constructed by replacing the *aac(3’)IV* gene with the *bla* or *neo* resistance genes, respectively, using In-Fusion cloning. All primers used for these clones are listed in Table 2.

### Construction of the pGGTOX-Cas9-sgRNA_KPC_ plasmid

The Cas9 encoding gene and sgRNA-N20_KPC_ sequence were amplified from the pCasCure-N20_KPC_ plasmid(21) using the primer pair ggTox-cureF/R. The pGGTOX plasmid was linearized by PCR using the primer pair cure-ggTOX-F/R. The amplified fragments were assembled into the linearized pGGTOX backbone via In-Fusion cloning to generate the recombinant plasmid pGGTOX-Cas9-sgRNA_KPC_. This construct enables targeted delivery of Cas9-sgRNA_KPC_ into recipient strains (21).

### Construction of the pCP20-oriT plasmid

The construction of pCP20-oriT followed a similar strategy to pGGTOX. Using pCP20 as a template, we amplified the *FLP* recombinase (*FLIP*) gene and the *λ*-repressor gene with primers 1088flip-F and 1088flip-R. Additionally, the chloramphenicol resistance gene and the temperature-sensitive *Rep101* replication origin were amplified using primers 1088rep-F and 1088rep-R. To incorporate hygromycin resistance, we amplified the *hyg* gene from pSGKp- hygromycin using primers hyg-F and hyg-R. Finally, the *oriT* sequence was amplified from the conjugative plasmid pdCas-J23119-RFP-oriT using primers 1211oriT-F and 1211oriT-R. These four fragments were assembled into a single construct using seamless cloning and transformed into *E.coli* WM3064, resulting in the successful construction of the pCP20-oriT plasmid.

### Preparation of competent *E. coli* WM3064 cells and transformation procedures

#### Chemical transformation

Chemically competent *Escherichia coli* WM3064 cells were prepared using the Takara Competent Cell Preparation Kit, following the manufacturer’s protocol. For transformation, 50–100 ng of plasmid DNA was gently mixed with 100 µL of competent cells and incubated on ice for 30 minutes to facilitate DNA uptake. Cells were then subjected to heat shock at 42 ℃ for 45 seconds. Immediately following heat shock, cells were transferred to 900 µL of LB broth supplemented with 0.3 mM DAPA. The recovery incubation was carried out at 37 ℃ with shaking for 1 hour. Transformed cells were subsequently plated on LB agar containing appropriate antibiotics and DAPA for selection of recombinant colonies.

#### Electroporation

Electrocompetent *E. coli* WM3064 cells were prepared using a glycerol wash protocol. Cultures were grown in LB medium supplemented with DAPA to an optical density at 600 nm (OD₆₀₀) of 0.4–0.6. Cells were harvested by centrifugation at 4 ℃ and washed three times with ice-cold 10% (v/v) glycerol to remove residual salts. The final pellet was resuspended in electroporation buffer (10% glycerol) to a final volume of 100 µL per transformation.

For electroporation, 50–100 ng of plasmid DNA was added to the competent cell suspension and transferred to a pre-chilled 0.2 cm electroporation cuvette. Electroporation was performed using a Bio-Rad Gene Pulser set to 1.8 kV, 25 µF, and 200 Ω. Immediately after pulsing, cells were recovered in 900 µL of SOC medium supplemented with DAPA and incubated at 37 ℃ with shaking for 1 hour. Recovered cells were plated on selective LB agar containing DAPA and appropriate antibiotics.

### Insertion of homology targeting regions to pGGTOX

The appropriate upstream and downstream sequences of the targeted genes were amplified in separate PCRs. Specifically, homologous regions were amplified with pGG-BsaI-dap1F, pGG-BsaI-dap1R, pGG-BsaI-dap2F, pGG-BsaI-dap2R (for Nissle 1917 *dapA* gene); pGG-rm-mrk1F, pGG-rm-mrk1R, pGG-rm-mrk2F, pGG-rm-mrk2R (for *mrkCD* genes); bsmB-traF1, bsmB-traR1, bsmB-traR3, bsmB-traF3 (for *tra* gene cassette knockout); BsaI-IS1R-2R, BsaI-IS1R-2F, BsaI-IS1R-1F, BsaI-IS1R-1R (for insertion of Cas9-sgRNA-N20_KPC_ encoding genes). After gel purification of the resulting PCR product, the products were mixed with pGGTOX plasmid (molar ratio of 5:1) and assembled by golden clone. Each pair of homologous fragments were used in a 20 µl reaction with 75 ng pGGTOX plasmid (2:1 molar ratio of insert:vector backbone), T4 DNA ligase buffer, 1 µl T4 DNA ligase and 1 µl BsaI (or BsmBI when needed). Temperature cycling: 20 min @ 37 ℃; 30-60 repeats of (5 min @ 37 ℃, 5 min @ 16 ℃); 5 min @ 60℃, hold @ 4 ℃. The assembly mix was used to transform chemically WM3064 strain, serving as the pGGTOX donor strain in conjugation. Colony PCR was performed on transformant green colonies on apramycin agar plate to confirm the appropriate assembly.

### Transference of pGGTOX to target strains by conjugation

*E.coli* WM3064 harboring correct homologous fragments were mixed with recipient strains. Bacterial cultures were grown for no more than 18 h prior to conjugation experiments. For each of the donor and recipient strain cultures, an adjusted volume of 500 µl at an OD600nm of 1.0 was transferred to separate sterile 1.5-ml microtube. Cells were centrifuged at 20,000× g for 1 min and washed once in 200-µl sterile LB. The cells were then resuspended in 2.5 µl. Donor and recipient strains were next mixed at a 2∼10:1 ratio. the mixture was then applied to 0.45-μm filter paper on an LB agar plate supplemented with DAPA and 2% glucose. Conjugation mixtures were incubated at 37℃ overnight, before being resuspended and diluted 1/10 serially in sterile PBS. The diluted mixture was spread onto Mueller–Hinton agar containing apramycin (50 mg/L) and cultured at 37℃ overnight. The white and green exconjugants were selected to confirm the homologous recombination by PCR respectively (Fig. S1).

### Isolation and identification of double-crossover mutants

Green fluorescent strains undergone successful first recombination were further subjected to a secondary recombination event. A single positive clone was inoculated into 5 mL LB medium containing 2% glucose, (but no selective antibiotic), then incubated at 37℃ with shaking at 220 rpm for 2.5 hours until OD_600_ reached 0.4–0.6. Bacterial cultures were centrifuged at 3500 rpm for 5 minutes, the supernatant was discarded, and the pellet was washed twice with M9 medium containing 2% rhamnose. The final bacterial suspension (100 µL) was spread onto M9 agar plates with 2% rhamnose and apramycin (50 mg/L), followed by incubation at 37℃ overnight. The following day, the white double-crossover colonies could be identified by the colony color. Colonies were further confirmed by colony PCR (Fig. S1). When *dapA* gene was targeted, the glucose LB media, M9 wash solution and M9 agar were supplemented with 0.3 mM DAPA.

### Antibiotic resistance gene removal

To facilitate the excision of apramycin resistance cassettes, the pCP20-oriT plasmid was introduced into recombinant strains from *E.coli* WM3064 via conjugation. This plasmid features temperature-sensitive replication and thermal induction of FLP recombinase, enabling site-specific recombination at FRT sites. Transformants were initially selected at 30 ℃ on media containing hygromycin and chloramphenicol. To induce FLP expression and promote recombination, selected colonies were subsequently purified non-selectively at 43 ℃, a temperature that inhibits plasmid replication and activates FLP synthesis. Successful excision of the resistance cassette was verified by PCR using primers flanking the targeted deletion site. Loss of the pCP20-oriT plasmid was confirmed by the absence of hygromycin and chloramphenicol resistance, indicating both recombination and plasmid curing were achieved.

### Biofilm formation assay

Biofilm formation by strains JNQH950, JNQH373, and their mutants was quantified using crystal violet staining(25). Overnight cultures were diluted 1:1,000 in LB broth, and 200 μl was added to U-bottom 96-well microtiter plates. Plates were incubated statically at 37 ℃ for 24 h. After incubation, wells were washed four times with distilled water to remove planktonic cells. Adherent biofilms were stained with 125 μl of 0.1% crystal violet for 10 min, then rinsed six times with distilled water. Stained biofilms were solubilized with 150 μl of 30% glacial acetic acid and incubated for 10 min at room temperature. Absorbance at 590 nm was measured using a microplate reader (Thermo Scientific, IE). Each sample was tested in triplicate, and experiments were repeated twice.

### Antibiotic susceptibility testing

Minimum inhibitory concentrations (MICs) of meropenem and imipenem were determined using the broth microdilution method. The results of the minimum inhibitory concentrations (MICs) were interpreted based on the Clinical and Laboratory Standards Institute (CLSI) M100-S32 guidelines. *E. coli* ATCC 25922 served as the quality control strain. Each assay was conducted in triplicate across two separate days to ensure reproducibility.

### Conjugation assay to evaluate plasmid transfer efficiency

To assess the transfer efficiency of the IncN/FII plasmids, conjugation experiments were performed using *E. intestinihominis* strain JNQH617 as the donor and *E. coli* J53 as the recipient. Overnight cultures were mixed (1:1) and applied to 0.45 μm filter paper, which was then placed on an LB agar plate. Transconjugants were selected on agar plates containing meropenem (0.5 µg/mL) and sodium azide (100 µg/mL). Successful plasmid transfer was confirmed by PCR amplification of the IncN replication gene. Conjugation frequency was calculated by dividing the number of verified transconjugants by the total number of recipient cells (15).

### Data availability

The GenBank accession numbers that correspond to the annotated sequences of the vectors used in this study will be released after publication, which are as follows: pGGB; pTOX8-GFP; pGGTOX (PV792960); pCP20-oriT; pdCas-J23119-RFP-oriT; pSGKp-hygromycin; pSZU941-GFP, pGGTOX-amp, pGGTOX-kana.

The complete sequences of the chromosomes and plasmids of the strains used in this study have been deposited in the GenBank database under the following accession numbers: JNQH950 (NZ_CP136146.1 to NZ_CP136148.1), JNQH617 (SAMN52855604) and JNQH373 (NZ_CP091978.1 to NZ_CP091980.1)

## Discussion

This study introduces pGGTOX, a self-eliminating plasmid platform designed for recombination-based genome editing across diverse *Enterobacteriaceae*. By integrating Golden Gate assembly with conjugative plasmid transfer, pGGTOX enables efficient and precise modification of both chromosomal and plasmid DNA in *E. coli*, *K. pneumoniae*, *E. intestinihominis*, and *Salmonella* strains. To enhance post-editing marker excision, we further engineered pCP20-oriT, a derivative of the widely used pCP20 backbone(14), which improves FLP-mediated recombination efficiency while maintaining full compatibility with the pGGTOX system. The versatility of pGGTOX was demonstrated through targeted disruption of representative chromosomal genes (*dapA*, *mrkCD*) and plasmid elements (*tra* cassette, Tn*6011*), as well as insertion of Cas9-sgRNA into a conjugative IncX1 plasmid. These applications underscore the platform’s adaptability for functional genomics and antimicrobial resistance studies.

A key observation was the differential efficiency between chromosomal and plasmid editing. Plasmid-based modifications in *S. enterica* and *E. intestinihominis* were achieved via single-step homologous recombination, whereas chromosomal edits in *E. coli* and *K. pneumoniae* required sequential recombination events. This disparity may be attributed to the multicopy nature of plasmids. Deletion events mediated by long direct repeats are predominantly driven by recombination between two plasmid molecules (26). Such intermolecular recombination enhances the likelihood of intramolecular deletions occurring during plasmid replication, thereby facilitating more efficient editing. Notably, integration of the pGGTOX plasmid into a target plasmid was observed to promote plasmid loss following induction of toxin expression, particularly when selective antibiotics were omitted from minimal M9 medium (data not shown). Although the efficiency of plasmid curing via this approach was relatively low, it nonetheless offers a potential strategy for selective plasmid curing.

Compared to CRISPR/Cas9-based systems such as pCasKP-pSGKP (5), the pGGTOX platform offers several distinct advantages. While CRISPR/Cas9 is highly effective for chromosomal editing, its application to plasmid targets often leads to unintended plasmid loss and carries the risk of off-target genomic alterations(21, 27, 28). Additionally, CRISPR-based approaches typically rely on labor-intensive delivery methods such as electroporation or chemical transformation, which can limit scalability and host range(5, 21). In contrast, pGGTOX facilitates direct conjugative transfer of editing constructs, eliminating the need for competent cell preparation and enabling broader accessibility across diverse bacterial species. Once a gene-targeting plasmid is constructed, it can be directly transferred via conjugation to edit the same genes in similar bacterial hosts, bypassing the need to prepare competent cells or perform transformation. From this point, a library based on pGGTOX tool targeting all the unessential genes of enterobacterols can be constructed for further convenient genome editing (29).

The system also supports large-scale genetic modifications, as demonstrated by the efficient deletion of a 43.1 kb *tra* cassette and insertion of a 10 kb Cas9-sgRNA module. This capability is likely supported by the robust homologous recombination machinery in *Enterobacteriaceae* (e.g., RecA, RecBCD, RecFOR), suggesting that even larger DNA fragments may be amenable to editing(30, 31). The inclusion of selectable antibiotic resistance markers upon gene disruption further streamlines mutant screening and functional analysis.

However, a limitation of the current system is the retention of an FRT site following FLP-mediated recombination. The residual FRT “scar“ remains at the target locus. This may pose challenges for downstream applications requiring scarless genome editing.

## Conclusion

In summary, we successfully developed and validated pGGTOX, a novel plasmid-based gene-editing tool that leverages homologous recombination for precise genomic modification. Designed for use across diverse *Enterobacteriaceae* strains, pGGTOX exhibited robust performance characterized by high editing efficiency, directional accuracy, and broad applicability. This system significantly enhances the existing microbial gene-editing toolkit by offering a streamlined platform for genetic manipulation. Its implementation facilitates more efficient functional genomics studies and accelerates the engineering of microbial strains for research and biotechnological applications.

---

**Fig. S1.**
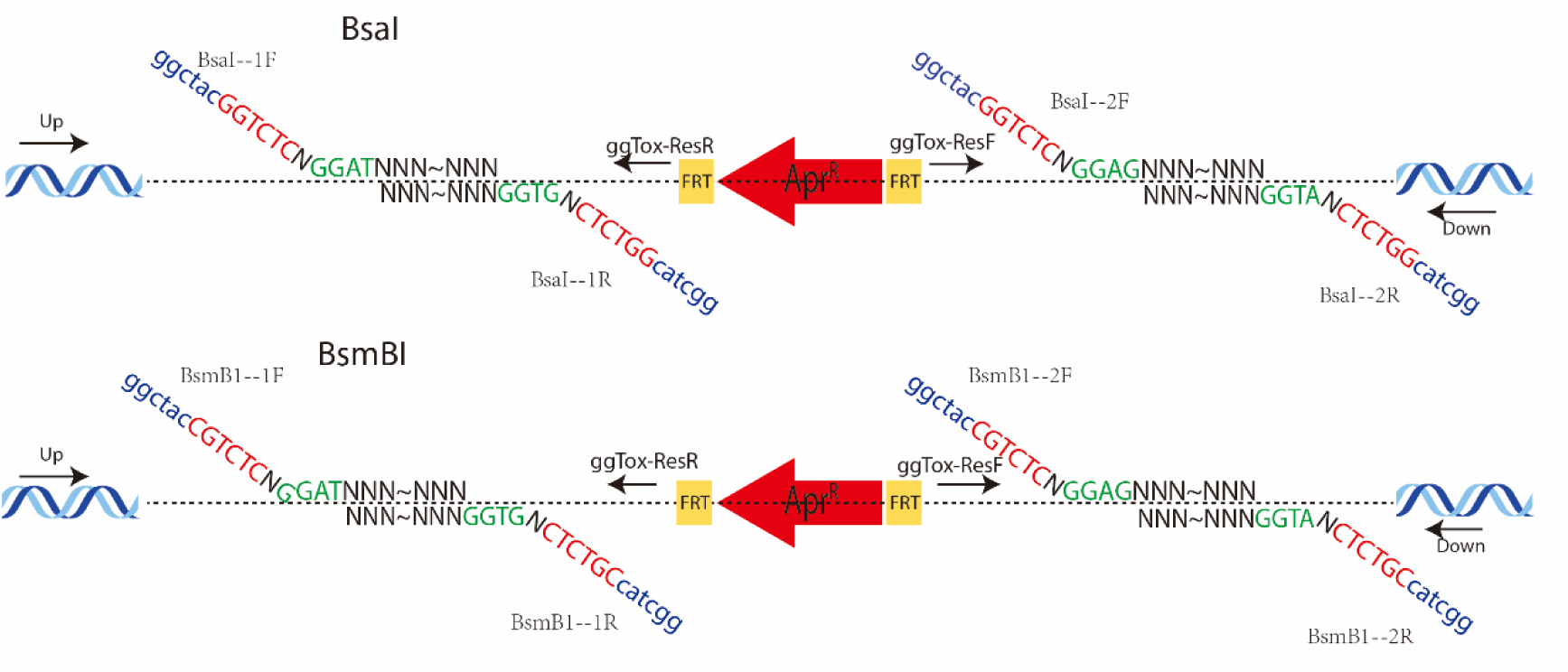
A PCR-based approach to evaluate the insertion of homologous fragments into the pGGTOX vector and to analyze the resulting gene editing outcomes. Homologous fragments (denoted as *N*) were amplified using primer pairs designed with type IIS restriction sites for directional cloning. Primer ends incorporated protection bases (blue lowercase), type IIS enzyme recognition sites for BsaI or BsmBI (red uppercase), and sticky 5′ overhangs generated upon enzymatic digestion (green uppercase). The primers *Up* and *Down* anneal to sequences flanking the homologous regions in the target gene. To confirm successful insertion of homologous fragments into the pGGTOX vector, PCR was performed using the primer pairs BsaI(BsmB1)-1F/ggTox-ResR and BsaI(BsmB1)-2R/ggTox-ResF. Following conjugation of pGGTOX variants carrying two homologous arms into the target strain, gene editing was assessed by PCR using Up/ggTox-ResR and Down/ggTox-ResF. A single positive PCR (either Up/ggTox-ResR or Down/ggTox-ResF) indicates recombination at one homologous arm. Dual positivity confirms recombination at both arms, signifying successful gene editing.

